# Selection on the joint actions of pairs leads to divergent adaptation and coadaptation of care-giving parents during pre-hatching care

**DOI:** 10.1101/2022.05.23.493134

**Authors:** Benjamin J. M. Jarrett, Rahia Mashoodh, Swastika Issar, Sonia Pascoal, Darren Rebar, Syuan-Jyun Sun, Matthew Schrader, Rebecca M. Kilner

## Abstract

The joint actions of animals in partnerships or social groups evolve under both natural selection, from the wider environment, and social selection, imposed by other members of the pair or group. We used experimental evolution to investigate how jointly expressed actions evolve upon exposure to a new environmental challenge. Our work focused on the evolution of carrion nest preparation by pairs of burying beetles *Nicrophorus vespilloides*, a joint activity undertaken by the pair but typically led by the male. In previous work, we found that carrion nest preparation evolved to be faster in experimental populations without post-hatching care (No Care lines) than with post-hatching care (Full Care lines). Here we investigate how this joint activity evolved. After 15 generations of experimental evolution, we created heterotypic pairs (No Care females with Full Care males, and No Care males with Full Care females) and compared their carrion nest making with homotypic No Care and Full Care pairs. We found that pairs with No Care males prepared the nest more rapidly than pairs with Full Care males, regardless of the female’s line of origin. This suggests that males led the way by adapting their nest preparation behaviour to the No Care environment first, with females secondarily co-adapting their behaviour to the male’s behaviour by reducing their nest preparation behaviour. We discuss how social coadaptations within pairs or groups could act as a post-mating barrier to gene flow.

## Introduction

When animals interact in partnerships, families or social groups, they are often in conflict, for example, over mating (Chapman et al. 2003), parental care (Royle et al. 2012), or who gets to reproduce (Emlen 1995). But, when the conflict is resolved or suppressed, their joint activities can enhance the fitness of the whole collective. Joint actions commonly function to overcome ecological adversity in the wider environment (e.g., Cornwallis et al. 2017), such as the threat of attack by predators (e.g., Feeney et al. 2013), the potential theft of a key resource by rivals (e.g., Queller & Strassman 1998) or the patchy availability of nutritional resources (e.g., Faulkes et al. 1997). It might involve the joint construction of a nest or communal burrow (e.g., Hansell 2005), for example, or collective defence against pathogens (e.g., Cremer 2019) or group care of dependent offspring (e.g., Queller & Strassman 1998). Working together to achieve a common goal favours individuals whose activities are well-coordinated with the actions of their partner, or other members of the family or social collective, (e.g., Barta et al. 2014), perhaps through division of labour (e.g., Cooper & West 2018). It means that for each individual in the collective, there are two inter-related sources of selection on their behaviour: 1) natural selection from ecological challenges in the wider environment that the team is working collectively to overcome and 2) social selection from the other members of the pair or group to fine-tune individual contributions to the pair or group’s collective activities (Itzkowitz et al. 2001, Barta et al. 2014, Cooper & West 2018, McNamara et al. 1999).

Theory predicts that adapting to the heritable environments created by social partners can be more rapid than adaptation to other environments. Selection imposed by the social environment changes the genetic variation within the heritable environment and reinforcing selection in the subsequent generations in a positive feedback loop (Moore, Brodie, and Wolf 1997, Agrawal, Brodie, and Wade 2001, Drown and Wade 2014). Furthermore, the social co-adaptation of the different group members establishes favourable combinations of phenotypes within the family. These are known to exist within animal families, for example (e.g., Zeh and Zeh 2000; Linksvayer, Fondrk, and Page Jr 2009; Hinde, Johnstone, and Kilner 2010), particularly among provisioning adults who coordinate their activities very precisely (e.g., Savage and Hinde 2019; Griffith 2019; Smiseth et al 2019). Favourable phenotypic combinations potentially result from intergenomic epistasis in which the fitness of a social trait depends on the genotype of the social partner (Wolf, Brodie, and Wade 2000; Wolf 2000).

The twin sources of selection on social traits, and the distinct patterns of inheritance and evolutionary dynamics of socially selected traits, make it difficult to predict how social traits might evolve upon exposure to a new ecological challenge in the wider environment. Would all individuals in the pair or group experience the same selection pressures in response to a new ecological challenge in the first instance, and only secondarily respond to selection induced by their behaviour to each other? Or would selection act on a subset of the population who first adapt to ecological change, and their new actions then provoke new social co-adaptations in other members of the pair or group? The latter scenario seems more likely for social activities in which there are pre-existing leaders and followers (e.g., Nagy et al. 2010), or where labour is already unevenly divided (e.g., Hager and Johnstone 2003; Hinde, Johnstone, and Kilner 2010; Kölliker, Royle, and Smiseth 2012; Barta et al. 2014), though whether this is indeed what happens has seldom been tested experimentally.

We investigated this problem by establishing lines of burying beetles *Nicrophorus vespilloides* and allowing them to evolve in environments with a distinct ecological challenge that we imposed experimentally. Burying beetles breed on the body of a small dead vertebrate (Scott 1998), commonly in pairs but sometimes as lone adults or multi-adult groups (Müller et al. 2007). The parents jointly convert the dead mouse or bird into a carrion nest by removing the fur or feathers, rolling it into a ball, covering it in anti-microbial anal exudates, and burying it. Eggs are laid in the soil surrounding the sunken carrion nest, and newly hatched larvae crawl to the nest where they aggregate. To help their larvae take up residence on the carrion nest, and feed upon it, parents may make an incision in the flesh prior to their arrival. The joint nest preparation activities of the parents are the focus of interest here. Parents also guard and feed larvae after they hatch, though larvae can survive in the laboratory with no post-hatching care at all (Eggert, Reinking, and Müller 1998).

In wild populations, pairs are under selection to prepare and bury the carrion nest as effectively and efficiently as possible. Carrion nest preparation is key for concealing this vital resource from rivals who might wish to use it for their own reproduction, including blowflies (Sun and Kilner 2020) and other carrion-breeding insects, congenerics and conspecifics and microbes (Scott 1998). We imposed selection on the pair’s joint carrion preparation activities by removing parents 53h after pairing, just before their offspring started to hatch (the ‘No Care’ lines). In two other experimentally evolving lines, parents were allowed to stay with their larvae throughout their development and so were able to provide both pre-hatching and post-hatching parental care (the ‘Full Care’ lines).

During the first twenty generations of experimental evolution, the No Care lines adapted rapidly to our experimental intervention (Schrader, Jarrett, and Kilner 2015a; Schrader et al. 2017), and exhibited divergent phenotypes in both larval (Jarrett et al. 2018a; Jarrett et al. 2018b; Rebar et al. 2020) and parental traits (Jarrett et al. 2018a; Duarte et al. 2021; Rebar et al. 2022). Of particular relevance here, No Care parents evolved to frontload their parental care. They became more efficient and more effective at converting carrion into a nest for their larvae, and in this way increased their offspring’s chance of surviving without post-hatching care (Duarte et al. 2021). In particular, No Care parents were more likely to make an incision into the carcass prior to larval hatching. This incision is noticeable upon inspecting the carcass and its presence prior to larval hatching strongly predicts brood success when larvae are deprived of any post-hatching care (Duarte et al. 2021; Eggert, Reinking, and Müller 1998; Jarrett et al. 2018a). Crucially, all the adaptations to a life without parental care that occurred within fifteen generations of experimental evolution were consistent between replicate populations, including the timing of making the feeding incision (Jarrett et al. 2018; Duarte et al. 2021).

In common with many other species (Henshaw et al. 2019), burying beetles divide the duties of parental care between the sexes. Males carry out more of the pre-hatching duties of care and take the lead in preparing the carrion nest (Walling et al. 2008; De Gasperin et al. 2016), while females are more involved with post-hatching care (Walling et al. 2008). Previous work on burying beetles suggests that task specialisation causes the sexes to be dependent upon each other, in the sense that they each perform more effectively when paired (Pilakouta et al. 2018). After fifteen generations of experimental evolution, we conducted crosses between the experimental lines to investigate whether No Care males alone had unilaterally adapted their carrion preparation in the No Care lines, or whether this was true of both sexes. We generated homotypic pairings (No Care adults paired with each other, and Full Care adults paired with each other) and heterotypic pairings (No Care adults paired with Full Care adults). With this experimental design, we were able to test the extent to which each sex had co-adapted its nest preparation behaviour to that of its partner. We also used these data to investigate whether the presence of a feeding incision predicted the survival of the brood when parents were present or absent after hatching.

We predicted that if the males alone unilaterally adapted in the No Care lines, then carrion prepared by No Care males should consistently bear a feeding incision with a higher probability than carrion prepared by Full Care males—regardless of the female’s line of origin. If the No Care females had co-adapted to the No Care males, then carrion prepared by homotypic No Care pairs should be more likely to carry an incision than carrion prepared by No Care females paired with Full Care males. By contrast, if the sexes had each unilaterally evolved changes in their nest preparation behaviour in the No Care lines, then the Full Care homotypic pairings should be least likely to bear a feeding incision 53h after pairing, while the No Care homotypic pairings should be most likely to have an incision at this point and both heterotypic pairings should lie somewhere between the two.

## Methods

### Experimental evolution

The experimentally evolving lines used in this work have been described in detail elsewhere (Schrader et al. 2017). In brief, we established a large founding population of 671 *N. vespilloides* individuals by interbreeding wild-caught individuals from four different woodlands in 32 pairs. Only one pair of the 32 comprised individuals from the same woodland population. Offspring from each of the 32 pairs were represented in each of the four replicate experimental lines. In two lines, larvae experienced Full Care (FC) in each generation, with both parents allowed to stay in the breeding box to feed and interact with their offspring. In the remaining No Care (NC) lines, both parents were removed from the breeding box at each generation, 53 h after they were paired, once carcass preparation was complete but before any larvae had hatched (see Schrader et al. 2017 for more details on the experimental set up). Pairs only bred once and only had one partner. FC lines had 30 pairs breeding per generation, and NC lines had 45 pairs to account for failures. Siblings and cousins were prevented from breeding. The lines were organised into two blocks, separated by a week, to ease the workload (hence NC1, FC1 in Block 1 and NC2, FC2 in Block 2). The experimental work reported here used lines that had been exposed to 15 generations of experimental evolution.

### Experimental design

The design of the experiment is shown in Figure 1. Prior to testing, the four lines went through a common garden environment of Full Care (where all parents were allowed to interact with their larvae throughout development). The aim was to standardise parental effects across all four treatments so that any residual variation could be attributed to evolutionary change. To set up the common garden generation, the FC1 and NC1 parents were bred three weeks after eclosing as adults, and the FC2 and NC2 parents were bred at two weeks after eclosing as adults. This synchronised breeding of individuals from the two blocks ensured that the adults generated for the focal experiment were matched in age at first breeding across treatments. The lines were then crossed: homotypic pairs involved crossing the two replicate lines that had experienced the same parental manipulation (i.e., FC1 × FC2 and NC1 × NC2, Fig 1a, b), while heterotypic pairs involved crossing populations that had experienced different parental manipulations (i.e., FC1 × NC1 and FC2 × NC2, Fig 1a, b). It was not feasible logistically to perform every possible cross between every replicate line (Fig 1a). With the experimental design we used, we can isolate the evolutionary effects of parental care treatment (NC v FC) on males and females (Fig 1a), but we cannot partition effects of replicate line (1 v 2) from parental care treatment (NC v FC). Note that when we carried out this experiment, we had detected no evidence of divergence between replicate lines with parental care treatments.

**Figure 1.**
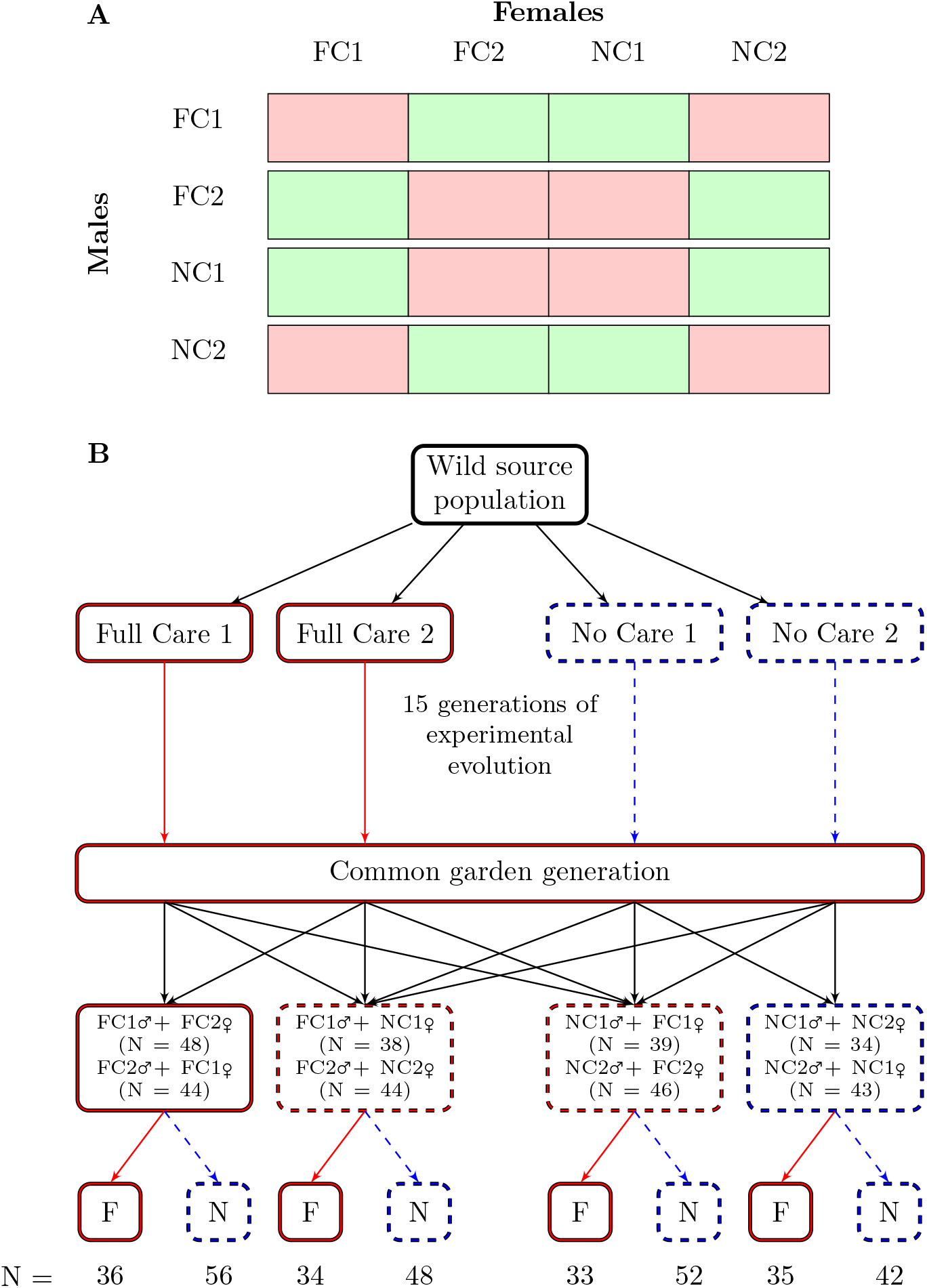
(A) Experimental design, showing all possible crosses between lines, including all replicate lines within parental care treatments. Green boxes indicate the crosses that were made, red boxes indicate the crosses that were not made. FC = Full Care, NC = No Care; numbers refer to replicate lines. (B) The experimental design in detail. Four lines were generated from a source population that formed two replicate Full Care (in red) and two replicate No Care (in blue) lines. The Full Cares lines evolved in a Full Care environment (red solid lines) and the No Care lines evolved in a No Care environment (blue dashed lines) for 15 generations (see Schrader et al. 2017 for more information about the experimental set up). After 15 generations of breeding in these contrasting environments, individuals from each population were passed through a common garden regime, in which all broodsreceived Full Care to minimise variation between lines due to parental effects. The populations were then crossed. The two control groups involved crossing the populations that had evolved under the same social environment during development: Full Care pairs (left red solid box) were formed by males and females from FC1 and FC2; and No Care pairs (right blue dashed box) were formed by males and females from NC1 and NC2. The two experimental groups (middle red dashed boxes) involved crossing populations that have evolved under different social environments during development. We only crossed one Full Care line with its No Care replicate (FC1 with NC1, and FC2 with NC2) for both sets of crosses. The offspring from each pair were then randomly allocated one of the social environments (Full Care or No Care) as a treatment during larval development.

### Burying beetle husbandry

Adult beetles were kept individually in boxes measuring 12 × 8 × 2 cm filled with compost for two weeks after eclosion until they were sexually mature. Individuals were fed ∼0.3 g of minced beef twice a week. Once individuals were sexually mature, they were paired for breeding. The pair of beetles was added to a larger box measuring 17 × 12 × 6 cm half filled with fresh compost. A mouse carcass, sourced from Live Foods Direct Ltd, was weighed and recorded (mean ± sd = 11.60 ± 0.87g; range = 10.00–13.18g) and placed into the box, after which the pair of beetles was added. The pair of beetles prepared the mouse carcass for the first two days, whilst the female laid eggs in the soil surrounding the carcass. At ∼53 h, when carcass preparation and egg laying had been completed, we removed the parents in the No Care treatment. For the Full Care treatment, the parents were left in the box until the larvae dispersed, typically eight days after pairing.

Eight days after pairing, the larvae had completely eaten the carcass and were ready to complete development. We removed the larvae, counted them, and weighed the whole brood to the nearest 0.0001g. After they had been weighed, the larvae were placed into an eclosion box, measuring 10 × 10 × 2 cm, with 25 individual cells, each 2 × 2 × 2 cm. An individual larva was placed in each cell and covered with peat that was sifted to remove large chunks of soil. Each box held one brood. Water was sprayed over the top to prevent desiccation during subsequent development, which typically lasted 18–21 days.

### Statistical analysis

All analyses were performed in R (version 3.5.1) (R Development Core Team 2018) using Bayesian models in Stan (Stan Development Team 2021) implemented in *brms* (Bürkner 2017; 2018). Additional packages used for the analysis and plotting of data in this paper include: *tidyverse* (Wickham 2017), *ggplot2* (Wickham 2009), *tidybayes* (Kay 2019), and *modelr* (Wickham 2018). Bayesian models run through *brms* provide an estimate of the slope (*β*) with 95% credible intervals (CI), which gives an indication as to the effect of the variable in question. If the 95% CIs for the *β* estimate did not overlap with 0, we assessed the model’s predictive ability by comparing the model containing the variable of interest with a simpler model that did not contain the variable (or interaction of variables) of interest. We compared these models using leave-one-out (loo) cross-validation and calculated the relative weight of each model to evaluate which model had the highest likelihood of predicting future data (Vehtari et al. 2017). Models fit in this manner move away from the dichotomy of deciding whether a variable is significant or non-significant and assess both the strength of a variable’s effect, and the predictive ability of the model. Post-hoc contrasts between different pairs were performed using the “hypothesis” function in *brms*.

We tested our predictions by focusing on the presence of a feeding incision on the prepared carrion nest at 53h after pairing, using data from both the No Care and Full Care post-hatching environments (see Fig. 1) since parents in both treatments had experienced the same opportunity to create a feeding incision by this point. (Note, that when we refer to the ‘No Care environment’ and the ‘Full Care environment’, we mean the within-generation treatment at the end of the experiment (shown in Fig. 1). We refer to the evolving lines as either the ‘Full Care lines’ or the ‘No Care lines’). We treated the presence or absence of a feeding incision on the carcass as a dummy variable with a Bernoulli distribution where 0 = no feeding incision and 1 = feeding incision present.

### The adaptive value of the feeding incision

We also investigated the fitness consequences of the feeding incision by asking how its presence or absence influenced the likelihood that a brood would succeed (i.e., that the brood would have at least one surviving larva at dispersal). We anticipated the relationship between the feeding incision and brood success would be dependent on the care environment, and so split the No Care and Full Care environment data and analysed each separately. For this analysis, we analysed brood success (success vs fail) assuming a Bernoulli distribution. The model included male line of origin (i.e., descended from a No Care or Full Care line) and female line of origin (i.e., descended from a No Care or Full Care line) and their interaction, along with the presence or absence of a feeding incision as a two-level factor.

### Adaptation and co-adaptation

We tested whether the lines of origin of both males and females each independently predicted the presence of a feeding incision on a carcass. We coded males and females from the Full Care lines as 1 and individuals from the No Care lines as 0. The model included both the female and male lines of origin and their interaction. Simpler models were then considered by first removing the interaction between female and male lines of origin, then secondly removing each line of origin in turn, and finally considering a model without either line of origin. For all analyses, the standardised (mean = 0, sd = 1) mass of the carcass, and the standardised size of the male and the female were included as covariates. Other covariates included in specific models are outlined below.

## Results

### The adaptive value of the feeding incision

In the No Care environment, the presence of a feeding incision before larval hatching increased the success of the brood (*β* = 1.02, [0.28, 1.91], Fig. 2). Brood success could not be further explained by any other divergence between the populations of origin, as the null model without female or male background had the lowest loo values and half the model weight (Table 1). However, in the Full Care environment, the presence of a feeding incision did not influence the success of the brood (*β* = 0.21, [-1.53, 2.27], Fig. 2). Brood success was partly explained by the female’s line of origin, but this model had no more explanatory power than the null model as shown by the small Δloo and almost equal loo weight (Table 1).

**Figure 2.**
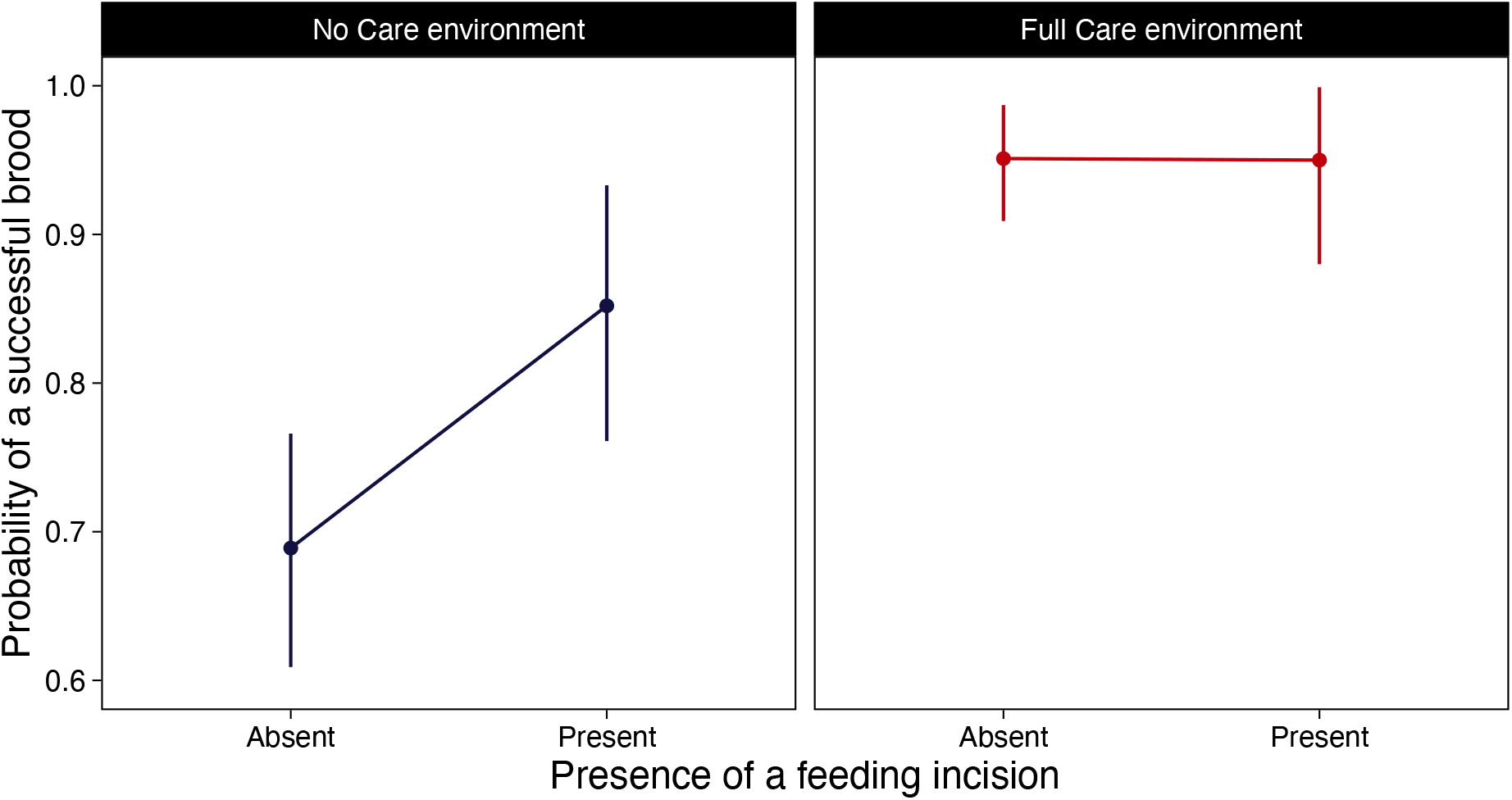
The proportion of successful broods when the feeding incision in the carcass was either present or absent prior to larval hatching, in a No Care and Full Care post-hatching environment. Predicted values (and 95% credible intervals) are shown derived from the model with carcass mass and female and male standardised size as other covariates.

**Table 1.**
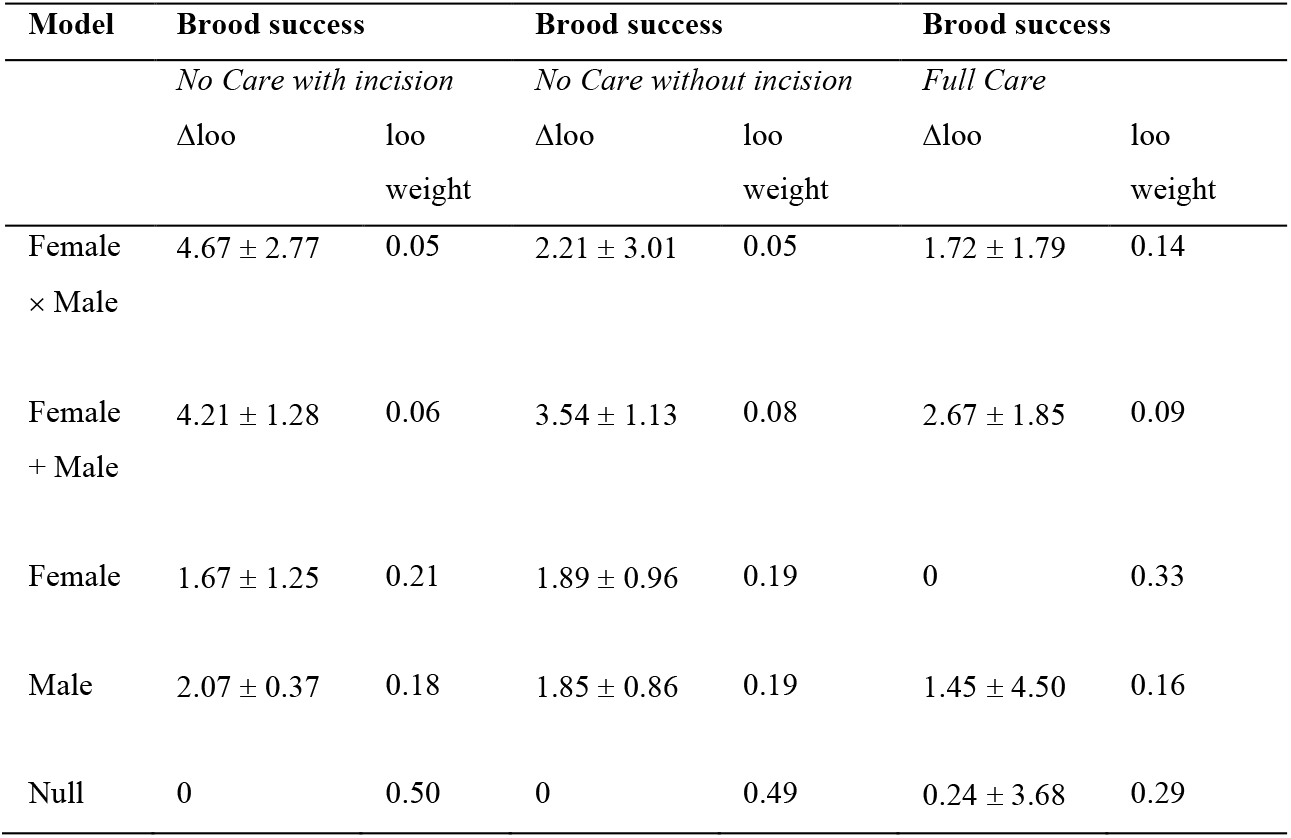
Summary of the model comparisons for brood success in No Care and Full Care environments, and, in the case of the No Care environment, whether the inclusion of the feeding incision as an explanatory variable alters which model best explains the data. A Δloo of 0 indicates it has the lowest looic value, with the difference and standard errors of the difference listed for each model. The looic weight is also listed for each model and sums to 1 for each set of models.

| Model | Brood success |  | Brood success |  | Brood success |  |
| --- | --- | --- | --- | --- | --- | --- |
|  | *No Care with incision* |  | *No Care without incision* |  | *Full Care* |  |
| | $\Delta\text{loo}$ | loo weight | $\Delta\text{loo}$ | loo weight | $\Delta\text{loo}$ | loo weight |
| Female<br>× Male | $4.67 \pm 2.77$ | 0.05 | $2.21 \pm 3.01$ | 0.05 | $1.72 \pm 1.79$ | 0.14 |
| Female<br>+ Male | $4.21 \pm 1.28$ | 0.06 | $3.54 \pm 1.13$ | 0.08 | $2.67 \pm 1.85$ | 0.09 |
| Female | $1.67 \pm 1.25$ | 0.21 | $1.89 \pm 0.96$ | 0.19 | 0 | 0.33 |
| Male | $2.07 \pm 0.37$ | 0.18 | $1.85 \pm 0.86$ | 0.19 | $1.45 \pm 4.50$ | 0.16 |
| Null | 0 | 0.50 | 0 | 0.49 | $0.24 \pm 3.68$ | 0.29 |

### Adaptation and co-adaptation

Replicating our previous finding (Jarrett et al. 2018a, Duarte et al. 2021), we found that homotypic No Care pairs were more likely to insert a feeding incision by 53h after pairing than homotypic Full Care pairs (Fig. 3, NN versus FF *post-hoc* comparison in Table 2).

**Figure 3.**
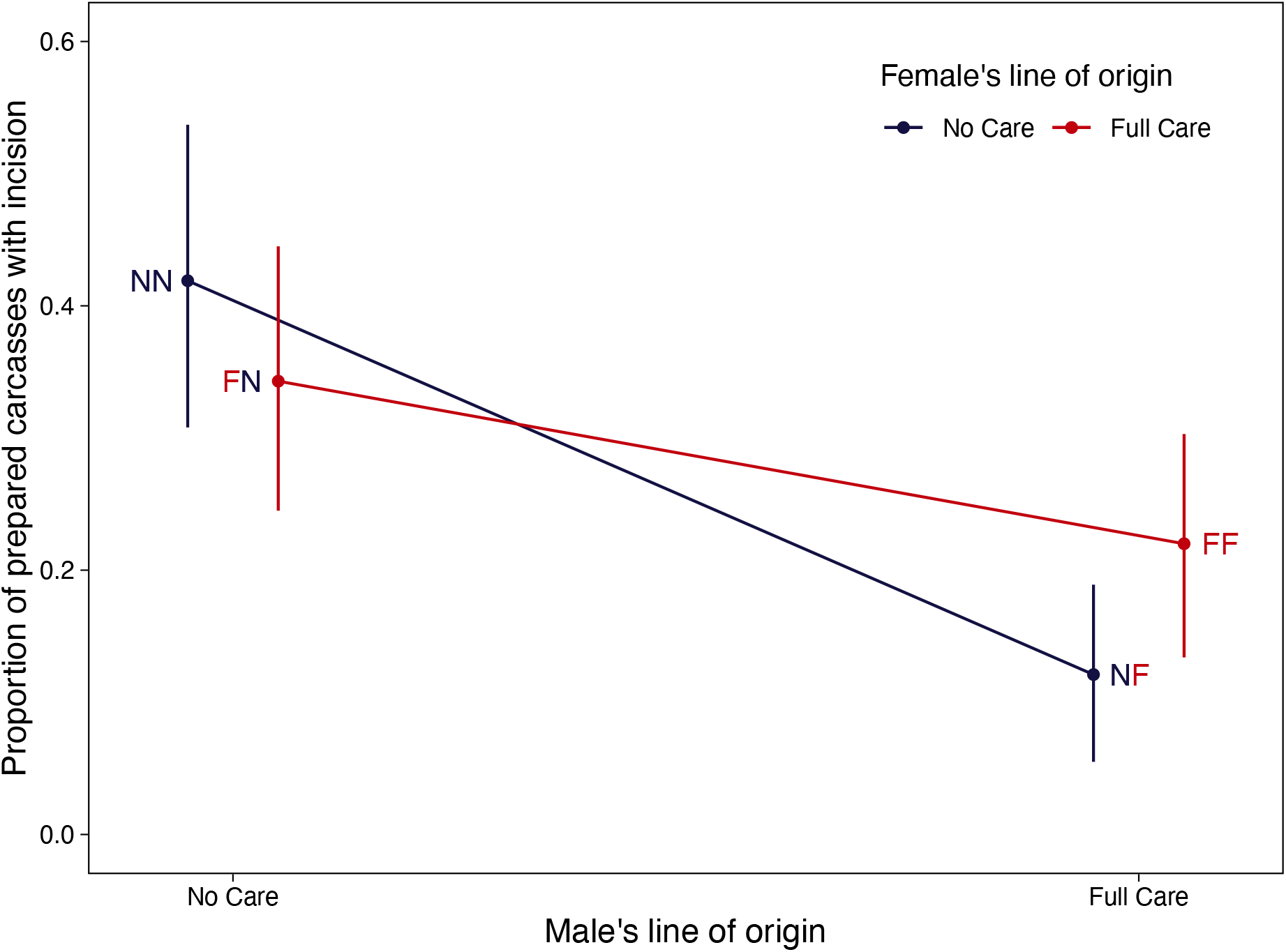
The proportion (and 95% credible intervals that describe the posterior distribution of the estimate) of prepared carrion nests found with an incision hole at 53 h after pairing, in relation to both the male and female’s line of origin. Predicted means are shown and are derived from the model containing the lines of origin of the male and the female and their interaction, as well as all other covariates.

**Table 2.** Post-hoc contrasts between all four combinations of female and male line of origin on the predicted probability of the presence of a feeding incision. The difference between the posterior distributions of the two pairs we are contrasting is shown, summarised by the mean and 90% CIs of the resulting distribution. Treatments are listed with female first and male second (for example, FN is a Full Care female paired with a No Care male). All comparisons are one-tailed save the FN vs NF comparison. We predicted that the NN would have greatest probability of making an incision and FF having the lowest probability, and so comparisons with these treatments were one-tailed accordingly. We had no *a priori* expectation for FN vs NF, which is why it is a two-tailed comparison. Credible intervals are 90% CIs except for the two-tailed test of FN vs NF which have 95% CIs. The final column identifies which of the two pairs in the contrast is most likely to make the incision, when the modulus of the contrast is greater than 0.

| Contrast | Estimate | Credible intervals | Pair more likely to make incision |
| --- | --- | --- | --- |
| FF vs NN | -0.95 | [-1.53, -0.36] | NN |
| FF vs NF | 0.74 | [0.05, 1.44] | FF |
| FF vs FN | -0.62 | [-1.18, -0.05] | FN |
| NN vs NF | -1.69 | [-2.38, -1.03] | NN |
| NN vs FN | -0.33 | [-0.86, 0.22] | neither |
| FN vs NF | -2.02 | [-3.21, -0.73] | FN |

Focusing first on males, we found that a feeding incision was more likely to be made if the male was drawn from a No Care population than a Full Care population (Fig. 3, Table 3). This was true regardless of the female’s line of origin. The No Care homotypic pair and the heterotypic pair of a No Care male and a Full Care female (Fig. 2, Table 3) did not differ in their likelihood of creating a feeding incision (Fig. 3; NN versus FN *post-hoc* comparison in Table 2). Therefore, we conclude that No Care males have unilaterally adapted their carcass preparation behaviour in response to the No Care treatment.

**Table 3.** Summary of the model comparisons the presence of the feeding incision. A Δloo of 0 indicates it has the lowest looic value, with the difference and standard errors of the difference listed for each model. The looic weight is also listed for each model and sums to 1 for each set of models.

| Model | Feeding hole |  |
| --- | --- | --- |
| | $\Delta loo$ | loo weight |
| Female<br>× Male | $0.11 \pm 4.21$ | 0.41 |
| Female<br>+ Male | $1.84 \pm 0.80$ | 0.17 |
| Female | $17.71 \pm 8.65$ | 0.00 |
| Male | 0 | 0.42 |
| Null | $15.56 \pm 8.46$ | 0.00 |

Turning to females, we found evidence that the presence of a feeding incision depended on the female’s line of origin matching the male’s line of origin (Fig. 3; interaction between sex and line of origin: *β* = 1.06, [–0.02, 2.15]). We accounted for the strong effect of a No Care male’s presence by focusing on comparisons with pairs involving Full Care males. Here, we found that a feeding incision was more likely to be made when a No Care female was paired with a No Care male, than with a Full Care male (NN vs NF comparison in Table 2). It was also more likely when a Full Care male was paired with a Full Care female rather than a No Care female (FF vs NF comparison in Table 2). Therefore, we conclude that females have co-adapted their carrion preparation behaviour to match the male’s behaviour.

## Discussion

We have previously shown that pairs of burying beetles evolved to prepare their carrion nest more rapidly when they were persistently prevented from interacting with their larvae after hatching (Duarte et al. 2021). Specifically, pairs evolved to hasten their insertion of a feeding incision in the carrion so that it was more likely to be present before their larvae hatched in the No Care lines than in the Full Care lines. The presence of a feeding incision contributes significantly to brood fitness when broods are deprived of any post-hatching care, because there is a greater chance that at least one larva survives until dispersal (Fig. 3; Duarte et al. 2021; Eggert, Reinking, and Müller 1998; Jarrett et al. 2018a).

We used this instance of experimental evolution to investigate a general problem. How do the joint actions of a pair (or group) evolve in response to change in the wider environment – in this case the enforced removal of parents prior to larval hatching? By setting up crosses within and between our experimental No Care and Full Care lines, we dissected the evolution of carrion preparation behaviour to determine how individual changes in males and females contributed to the accelerated expression of this pair-level trait in the No Care lines. We found that No Care males had unilaterally evolved to ensure the feeding incision was inserted sooner in the absence of post-hatching care. Nests prepared by No Care males were more likely to bear a feeding incision, regardless of the female’s line of origin. The likelihood that a feeding incision would be made prior to larval hatching was secondarily influenced by females, and the extent to which females were co-adapted to the behaviour of the male. Although both sexes contribute to carrion preparation, males take the lead in carrying out this duty of care. This might explain why evolutionary change in this pair-level trait was driven by the males in our experiments, with secondary co-adaptation by females, rather than *vice versa*.

The strength of our conclusions is tempered by the fact that our experimental design lacked any control pairings within replicate lines. (We had to sacrifice this treatment due to logistical constraints). Nevertheless, from other data we have collected from these populations, we have no reason to suppose that the replicate lines, within either the No Care or Full Care treatments, had diverged by this stage of experimental evolution (Schrader, Jarrett, and Kilner 2015a; Schrader et al. 2017; Jarrett et al. 2018a; Jarrett et al. 2018b; Rebar et al. 2020; Duarte et al. 2021). If our conclusions are broadly correct, then our expectation is that the evolution of other group or pair-level traits in response to environmental change should be initiated unilaterally by the individual that takes that lead in shaping these collective actions, and that this in turn will provoke swift social co-adaptation by other members of the pair or group.

The coadaptation of male and female traits is likely to have favoured a distinct combination of socially interacting genes in the two sexes (*cf* Linksvayer, Fondrk, and Page Jr 2009). For this reason, previous theoretical analyses have suggested that socially coadapted traits within the family that are divergent between populations could function as a post-mating mechanism for reproductive isolation—because they cause hybrids to perform less well (e.g., Zeh and Zeh 2000; Gavrilets 2000; Martin & Hosken 2003; Brandvain & Haig 2005). Evidence from burying beetles is consistent with this view. *Nicrophorus vespilloides* has recently been split into two species, with *N. hebes* now recognised as a distinct bog-breeding specialist that lives mostly in Canada (Sikes et al. 2016).

Speciation is sufficiently recent that *N. hebes* can still hybridise with Alaskan populations of *N. vespilloides* to produce viable offspring. Nevertheless, hybrids perform less well, partly because hybrid larvae are less viable (at least in a No Care post-hatching environment). Experimental work indicates that hybrid pairings produce fewer eggs and have lighter broods than pure-bred populations (Sikes et al. 2016). The reduced performance of hybrids could also be due to coadaptation between care-giving adults, although this possibility remains to be tested explicitly.

Social coadaptations contribute to post-mating reproductive isolation in other species (e.g., Zeh and Zeh 2000; Gavrilets 2000; Martin & Hosken 2003; Brandvain & Haig 2005). The key barrier is the high level of coordination between interacting individuals. This is true whether coordination results from cooperation or conflict. It is seen in viviparous species, for example, through the highly specific structures that have evolved to coordinate the supply of resources from mother to offspring in viviparous fish (Schrader & Travis 2008; Furness et al. 2019) and placental mammals (Roy 2022).

We have shown here that the highly coordinated activities jointly undertaken by male and female burying beetles in converting carrion into an edible nest can also become coadapted and cause heterotypic pairings to perform less well. In future work, it would be interesting to test whether the coadaptations involved in other types of coordinated cooperative social activities, such as the construction of nests or burrows, collective immunity, or the joint defence of a key resource, could also potentially function as a post-mating barrier to gene flow between populations.

## Acknowledgements

We thank Simon Martin, Jenny York, the Miller Lab, and Judith Mank for constructive comments and discussions. We are extremely grateful to Ercol Akcay, Tim Linksvayer and two anonymous reviewers for their helpful comments on previous versions of this paper. This project was supported by the European Research Council (310785 BALDWINIAN_BEETLES) and a Royal Society Wolfson Merit Award, both to R.M.K. Writing the first draft of this manuscript was supported by a Human Frontiers Science Program fellowship (LT000879/2020) to B.J.M.J. All data and R code is available at github.com/bjmjarrett/ghostbusters.

## Notes

### Competing Interest Statement

The authors have declared no competing interest.

### Summary of Updates

Introduction has been rewritten to deemphasise multilevel selection and intergenomic epistasis; removed clutch size data; improved Figure 1 for clarity of the limitations and strengths of the dataset; added post-hoc comparisons of the treatments.

